# Tumor extracellular vesicles are required for tumor-associated macrophage programming

**DOI:** 10.1101/375022

**Authors:** Daniel C Rabe, Felicia D Rustandy, Jiyoung Lee, Marsha Rich Rosner

**Author notes:** Corresponding author: Marsha Rich Rosner.

## Abstract

Triple-negative breast cancers (TNBC) are highly infiltrated by tumor-associated macrophages (TAMs) that promote tumor growth, survival, metastasis and therapeutic resistance. Although cytokines such as CCL5 have been implicated in TAM recruitment to TNBC tumors, the mechanism by which tumor cells educate TAMs is not understood. Here we show that tumor EVs are both necessary and sufficient for programming TAMs toward a pro-metastatic phenotype. The mechanism involves CCL5 regulation of tumor extracellular vesicles (EVs), which activate TLR2 and TLR3, leading to secretion of a common set of cytokines that further stimulate tumor cell invasion and metastasis as well as alter the tumor microenvironment. Cytokine expression is significantly correlated to CCL5 expression and up-regulated in TNBC patient tumors. These results demonstrate for the first time that tumor EVs are key mediators of TAM education, phenocopy the pro-metastatic and drug resistant state of the tumors to TAMs, and illustrate the potential clinical relevance of these findings to TNBC patients.

**Highlights:**

- Tumor extracellular vesicles (EVs) are required for pro-metastatic programming of tumor-associated macrophages (TAMs)
- Tumor CCL5 and macrophage TLR signaling mediate tumor EV programming of TAMs in TNBCs
- Tumor EVs mediate drug resistance in TAMs and alter recruitment of regulatory T-cells.
- Cytokines expressed by EV-educated TAMs are enriched and correlate with CCL5 in human TNBC patients.

**eTOC:** Chemokines such as CCL5 recruit tumor-associated macrophages (TAMs) that are required for metastasis, but TAM programming is not understood. Rabe et al. show that tumor extracellular vesicles (EVs) are required for programming TAMs via Toll-like Receptors (TLRs) to phenocopy the tumor, rewire the microenvironment, drive metastasis and promote immune cell evasion.

## Introduction

Of the women in the United States that develop breast cancer, 15-20% will have basal-like or triple-negative breast cancer (TNBC). Triple-negative tumors lack expression of estrogen receptor (ER), progesterone receptor (PR) and amplification of human epidermal receptor 2 (HER2) (Carey et al., 2006). Because of the poor response to chemotherapy and lack of currently approved targeted therapies for TNBC patients, their 5 year survival rate is only 24%; by contrast, overall 5-year survival rates are 89.7% (Howlader et al., 2013). Thus, it is important to identify mechanisms that drive metastatic progression of TNBCs and potential targets for therapeutic treatment.

Tumor cells are driven to metastasize in part through interaction with cells in the microenvironment. Tumor-associated macrophages (TAMs) are generally characterized as alternatively activated or M2 and secrete growth factors leading to tumor growth, progression, and metastasis (Green et al., 2009; Joyce and Pollard, 2009). M2 macrophages, thought to be similar in phenotype to TAMs, are commonly generated in culture from bone marrow-derived macrophages following programming with IL-4, IL-10, IL-13 or TGF-beta (Wynn et al., 2013). Recent work has shown that the level of M2 macrophages in the tumor stroma (determined through cluster of differentiation 163 (CD163) staining) correlates with outcome in patients, as well as with tumor type. In particular, M2 macrophage recruitment positively correlates with TNBC tumors while negatively correlating with ER+ tumors (Medrek et al., 2012). Therefore, recruitment of alternatively activated M2 macrophages could play a significant role in the outcome of TNBC patients and explain their poor prognosis. Work from Levano et al, 2011 has shown that basal versus luminal subtypes of breast cancer express different repertoires of cytokine receptors allowing them to be differentially affected by M2 macrophages (Levano et al., 2011).

Pollard and coworkers first examined the role of TAMs in breast cancer by knocking out macrophage colony stimulating factor-1 (CSF-1) in transgenic mice expressing the Polyoma Virus middle T antigen (PyMT) oncogene under the control of the mouse mammary tumor virus (MMTV) long terminal repeat (LTR). Although development of mammary tumors in these mice was unchanged, progression of the tumors to an invasive and metastatic state was delayed (Lin et al., 2001). To study the effect of TAMs on tumorigenesis, clodronate-tagged liposomes have been utilized to specifically ablate macrophages after tumor formation. Macrophage ablation reduced the ability of tumor cells to invade and metastasize (Joyce and Pollard, 2009) as well as potentiated the anti-tumor effects of targeted therapies in liver cancer (Zhang et al., 2010).

TAMs are recruited to mammary tumors through induction of a variety of cytokines and chemokines, where they play essential roles in driving metastasis. Much of the work in the immune environment of breast cancer has been done utilizing the MMTV-PyMT model of breast cancer. This model established CSF-1 and CCL2 as critical factors for recruiting TAMs to the primary tumor as well as metastatic sites. TAMs recruited by CSF-1 express higher levels of VEGF-A and promote increased angiogenesis in the MMTV-PyMT genetically engineered mouse model for breast cancer (Lin et al., 2006). Similarly, CCL2 was required for TAM infiltration into primary breast tumors as well as TAM-enabled metastatic colonization of lungs (Qian et al., 2011). When GFP monocytes were injected into the blood stream of mice along with tumor cells derived from the MMTV-PyMT model, a CCL2 neutralizing antibody blocked their recruitment to metastatic sites in the lung and subsequent metastatic outgrowth of tumor cells.

Work in other models of breast cancer has suggested that CCL5 plays a critical role in the recruitment of TAMs and other immune cells to tumors. Antagonists of CCL5 inhibited TAM recruitment in a syngeneic 4T1 mouse model (Robinson et al., 2003). CCL5 has also been implicated in the recruitment and activity of T cells. In particular, work has shown that tumor-derived CCL5 can recruit T-regulatory cells (T-regs) to the tumor, leading to CD8+ T-cell apoptosis (Chang et al., 2012). Blockade of CCL5 only decreased primary tumor growth in immune competent models, suggesting that T-cells are required to reduce tumor bulk.

Our previous work demonstrated that the metastasis suppressor Raf kinase inhibitory protein (RKIP) inhibits recruitment of tumor-associated macrophages (TAMs) to TNBC tumors by reducing CCL5 expression (Frankenberger et al., 2015). Overexpression of CCL5 in RKIP-transfected tumor cells rescued TAM infiltration. CCL5-recruited TAMs had increased expression of growth factors, cytokines, and other pro-metastatic factors including CCL7, MMP-12, and Progranulin, leading to an increased ability to drive tumor invasion *in vitro* and an increase in tumor intravasation *in vivo*. We generated a gene signature based on CCL5 signaling in tumors and TAM gene expression that specifically identified high-risk TNBC patients within a breast cancer cohort. Although this study established CCL5 as the dominant mechanism for recruiting TAMs to TNBC xenografts, the mechanism by which the macrophages are programmed or educated to a pro-metastatic M2-like phenotype is unclear.

A number of studies suggest that extracellular vesicles (EVs), including exosomes, modulate tumor, stroma, and immune response within the tumor microenvironment. Tumor-derived EVs have been implicated in the creation of a pre-metastatic niche (Costa-Silva et al., 2015; Hoshino et al., 2015) and in driving progenitor cells within the bone marrow toward a prometastatic state via MET expression (Peinado et al., 2012). EVs secreted by the stroma promote migration (Luga et al., 2012) and mediate resistance of breast cancer cells to chemotherapy (Boelens et al., 2014). Tumor-derived EVs are reported to stimulate myofibroblast differentiation (Webber et al., 2010) and to suppress immune response by blocking T cell activation by IL-2 (Clayton, 2012), inhibiting NK function (Liu et al., 2006) or potentiating regulatory T cell (T-reg) activity (Szajnik et al., 2010). EVs are also taken up by macrophages, likely through a phagocytic pathway (Costa-Silva et al., 2015; Feng et al., 2010; Kapsogeorgou et al., 2005). Furthermore, tumor EVs can stimulate cytokine production in mouse and human macrophage cell lines *in vitro* through activation of Toll-like receptors (TLR2-4) (Alexopoulou et al., 2001; Bretz et al., 2013; Chow et al., 2014; Liu et al., 2016) or through transfer of mRNAs transcribed in target cells (Skog et al., 2008). However, direct evidence that EVs direct tumor-associated macrophage education or programming is lacking.

Here we show that tumor EVs are both necessary and sufficient for the programming of macrophages to an M2-like, TAM phenotype in TNBC. Our results indicate that macrophages programmed by EVs are capable of driving invasion and metastasis as well as rewiring the tumor microenvironment toward a pro-tumor phenotype. Furthermore, we show that, while CCL5 can directly recruit macrophages to TNBC tumors (Frankenberger et al., 2015; Keophiphath et al., 2010; Robinson et al., 2003), macrophage reprogramming to TAMs occurs through an indirect mechanism involving autocrine stimulation of tumor cells by CCL5. This leads to EV release and subsequent macrophage reprogramming through a mechanism involving Toll-like receptors 2 and 3 (TLR2 and 3). We show that cytokines released by these tumor EV-educated macrophages (TEMs) correspond to cytokines induced in TAMs by tumor-expressed CCL5, and that TEMs injected with tumor cells into mice promote both TNBC tumor growth and metastasis to the lungs. Finally, manipulation of CCL5 levels in tumor cells through genetic or pharmaceutical alteration results in altered EV cargoes and altered TEMs that reflect the phenotype of the secreting tumor cells. Taken together, these results indicate that EVs are a key mechanism by which tumor-recruited macrophages are reprogrammed within the tumor microenvironment to a pro-metastatic TAM phenotype in TNBC.

## Results

### CCL5-recruited TAMs increase invasiveness in a persistent manner

Our previous work demonstrated that exogenous CCL5 expression in tumors expressing the metastasis suppressor RKIP was sufficient to recruit TAMs and restore TNBC tumor cell intravasation (Frankenberger et al., 2015). The tumor CCL5 signaling pathway, when combined with CCL5-induced TAM gene expression, formed the basis of a clinically prognostic gene signature we used for TNBC patients (Frankenberger et al., 2015). To directly assess the effect of CCL5-recruited TAMs on metastasis, we isolated TAMs from RKIP expressing (non-metastatic) and RKIP+CCL5 expressing (CCL5-rescued) tumors. We then injected one million MDA-MB-231 cells along with 0.5 million TAMs into nude mice. To address the effect on metastasis in this xenograft model, we assayed the ability of tumor cells to intravasate into the blood stream by comparing tumor *GAPDH* to mouse *Gapdh* expression using qRT-PCR as previously described (Frankenberger et al., 2015; Sun et al., 2013; Yun et al., 2011). We found that TAMs that were recruited and educated by tumor CCL5 expression showed an increase in both tumor growth (**Fig. 1A**) as well as tumor cell intravasation into the blood stream (**Fig. 1B**). As a negative control one group received TAMs from RKIP over-expressing tumors, which were not able to potentiate tumor growth or intravasation (**Fig. 1A-B**). These results suggest that TAMs recruited by CCL5 are programmed by tumor cells to regulate the metastatic properties of tumor cells.

**Figure 1:**
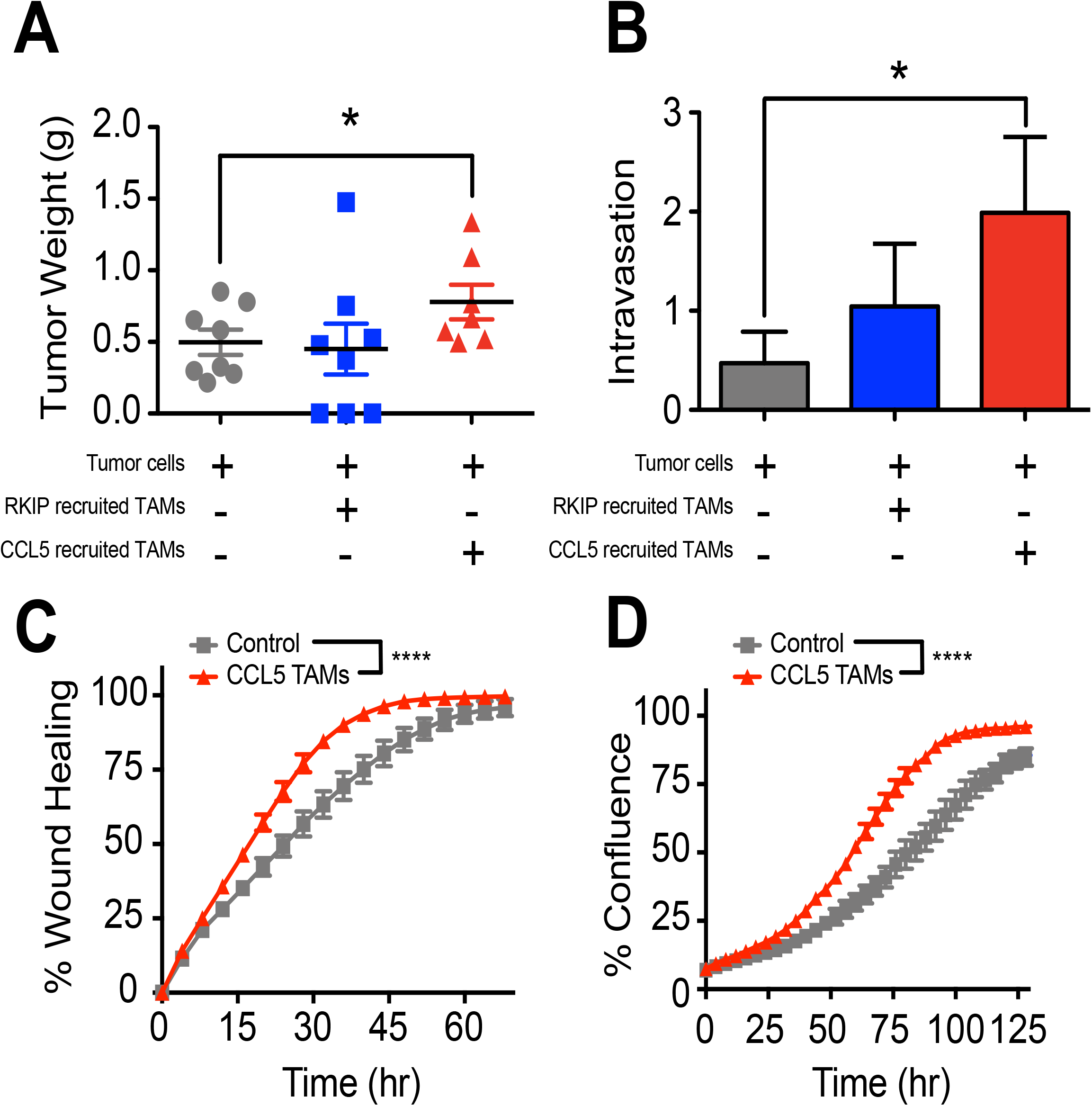
CCL5-recruited TAMs increase invasiveness in a persistent manner. MDA-MB-231 tumor cells (1 × 10^6^) were injected with or without TAMs (0.5 × 10^6^) into each mouse as indicated. TAMs were isolated from tumors that either over-expressed RKIP (RKIP TAMs, blue) or RKIP and CCL5 (CCL5 TAMs, red). **A)** Final tumor weights are shown per group. **B)** Relative intravasation was measured by quantifying the ratio of human *GAPDH* (tumor) to mouse *Gapdh* by qRT-PCR from blood taken immediately prior to sacrifice. P-values were calculated using a Student’s t-test (N=10). **C-D)** % Wound healing (**C)** and % Confluence **(D)** shown for MDA-MB-231 grown *ex vivo* from either Control (grey) or tumors co-injected with CCL5-recruited TAMs (red). P-values were calculated using a paired t-test (N = 8).

We next addressed whether the phenotypic alterations of tumor cells by TAMs were transient or persistent. To answer this question, we isolated cells from MDA-MB-231 tumors co-injected with TAMs and grew them in culture for 4 weeks. The tumor cells derived from mice that were co-injected with CCL5-recruited TAMs showed an increased growth rate (**Fig. 1D**) as well as an increased migratory ability compared to cells isolated from control tumors (**Fig. 1C**).

This suggests that changes in tumors cells resulting from co-injected TAMs are persistent, and do not require the prolonged presence of TAMs to be maintained.

### Tumor conditioned media programs macrophages toward a pro-invasive phenotype, and is altered by CCL5 signaling

To address how tumor cells program macrophages toward a pro-tumor TAM phenotype, we developed an assay based upon the ability of tumor cell conditioned media (CM) to educate bone marrow-derived macrophages (BMDMs). We termed these tumor-educated macrophages (TEMs) (**Fig. 2A**). We first asked whether the CM of the tumor cells is sufficient to alter gene transcription in BMDMs and generate TEMs that promote tumor cell invasion. TNBC CM was initially collected and used to program TEMs that were assayed for changes in expression of *Ccl7* (**Fig. 2B**), a gene induced in TAMs by CCL5-expressing tumor cells (Frankenberger et al., 2015) Ccl7 is known to bind CCR1, CCR2, and CCR3, which have been implicated in TAM recruitment and promotion of metastasis (Jung et al., 2010; Mishra et al., 2011). CM from the programmed TEMs was subsequently collected and used to pre-treat TNBC cells which were then assayed for invasion. TEMs were programmed using CM from BM1, the bone-metastatic variant of the human TNBC cell line MDA-MB-231, or LMB, the mouse TNBC cell line E0771-LMB. The results indicate that expression of *Ccl7* is induced in TEMs, and CM from the TEMs increases tumor cell invasion (**Fig. 2B**).

**Figure 2:**
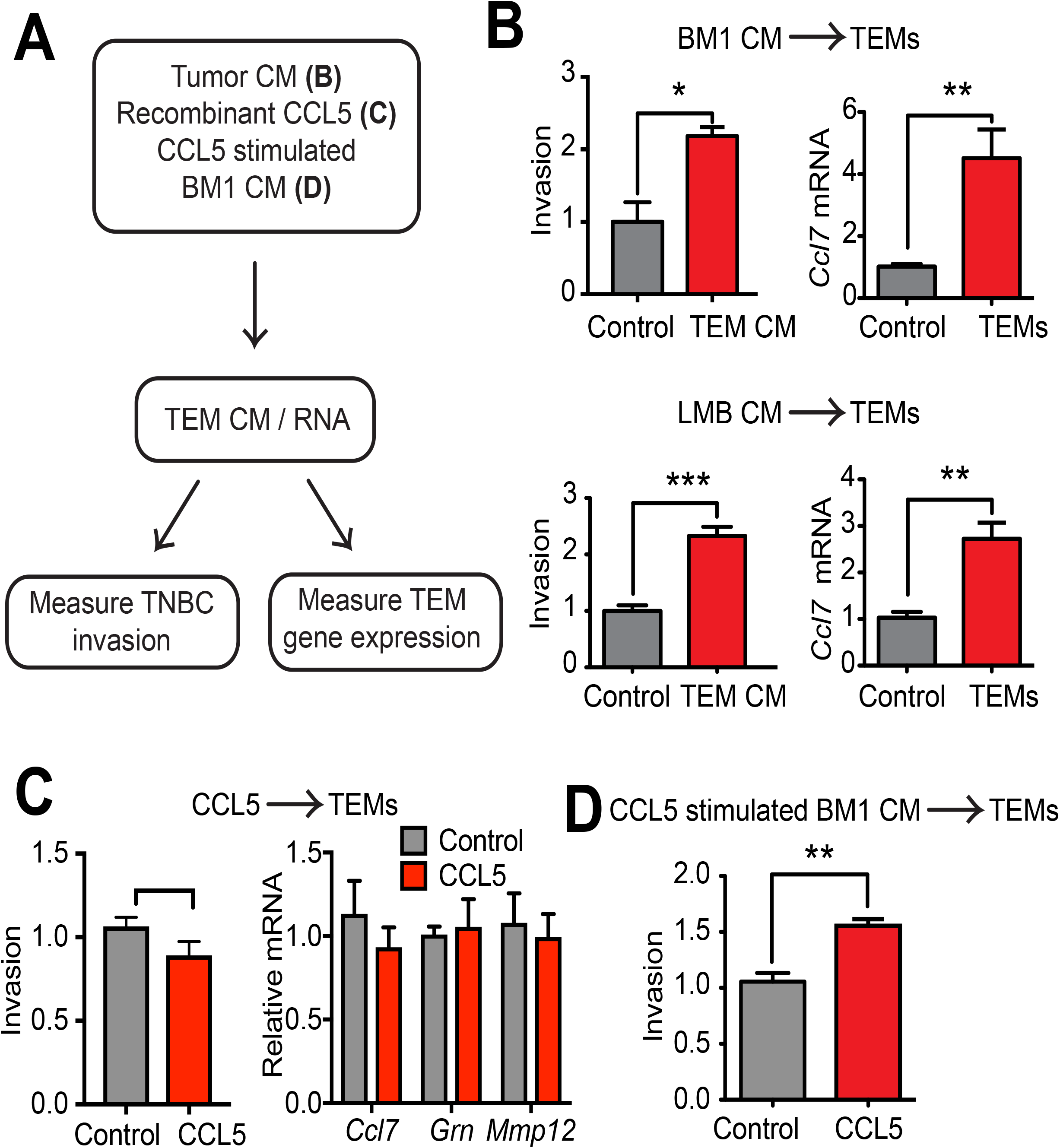
Tumor conditioned media programs macrophages toward a pro-invasive phenotype, and is altered by CCL5 signaling. **A)** Schematic depicting TEM programming and functional assays. **B**) Left: Relative invasion of TNBC cells pre-incubated for 24 hrs with TEM CM. Right: qRT-PCR of *Ccl7* in TEMs programmed with TNBC CM compared to M-CSF alone (Control) with *Gapdh* as a loading control. P-values were calculated using a Student’s t-test (N = 5). **C)** Invasion of TNBC cells pre-incubated for 24 hrs with TEM CM. **C)** Left: Relative invasion of BM1 cells pre-treated with either 20 ng/ml of M-CSF (Control) or 20 ng/ml of M-CSF plus 1 μg/ml of CCL5 for 48 hrs. Right: Relative expression in TEMs is shown for *Ccl7*, *Grn*, and *Mmp12* as measured by qRT-PCR, with *Gapdh* as the loading reference. **D)** Relative invasion of BM1 cells pre-incubated for 24 hrs. with TEM CM. TEMs were programmed for 48 hrs. with CM from BM1 cells treated for 24 hrs. with 100 ng/ml of CCL5 or serum-free media. P-values were calculated using a Student’s t-test (N = 5).

Since CCL5 recruits macrophage to tumors, we determined whether CCL5 protein could directly program TEMs by altering gene expression. Our previous work demonstrated that, when BMDMs were treated with a high dose of recombinant human CCL5 (1 μg/ml) compared to a BMDM control under serum-free conditions, *Grn*, *Mmp12*, and *Ccl7* expression were all induced (Frankenberger et al., 2015). However, when 20 ng/ml of mouse M-CSF is added to support macrophage survival in culture during TEM programming, additional stimulation of macrophages even in the presence of this high CCL5 dose was not sufficient to induce the expression of the pro-metastatic genes *Ccl7*, *Grn*, or *Mmp12* in the TEMs nor to program TEMs to a pro-invasive phenotype (**Fig. 2C**). These results suggest that M-CSF alone is capable of stimulating low-level expression of many pro-invasive genes, and addition of CCL5 in the presence of M-CSF is not sufficient to further induce expression above these levels.

Because CCL5 was not able to act directly on macrophages to program them, we next determined if CCL5 could act indirectly by stimulating tumor cells to secrete factors that program TEMs. BM1 cells were treated with serum free media as control or serum free media with 100 ng/ml of recombinant human CCL5 for 24 hours. After 24 hours, cells were washed to remove CCL5, and CM was collected over 24 hours to assay its ability to program TEMs. When we tested the CM from TEMs programmed in response to CCL5-stimulated BM1, we saw an increased induction of tumor cell invasion compared to control (**Fig. 2D**).

### EV programming of TEMs reflects the invasive state of the tumor and is regulated by CCL5

Previous work established that tumor EVs could regulate cytokine expression in macrophages *in vitro* (Bretz et al., 2013; Chow et al., 2014; Liu et al., 2016). We therefore determined if EVs within the CM are responsible for TEM programming **(Fig. S1)**. Upon centrifugation at 100,000 x g for 1 hr to deplete EVs, the CM of TNBC cells was no longer able to program TEMs to potentiate TNBC invasion (**Fig. 3A**). These results indicate that soluble factors in TNBC CM are not mediating the programming of TEMs.

**Figure 3:**
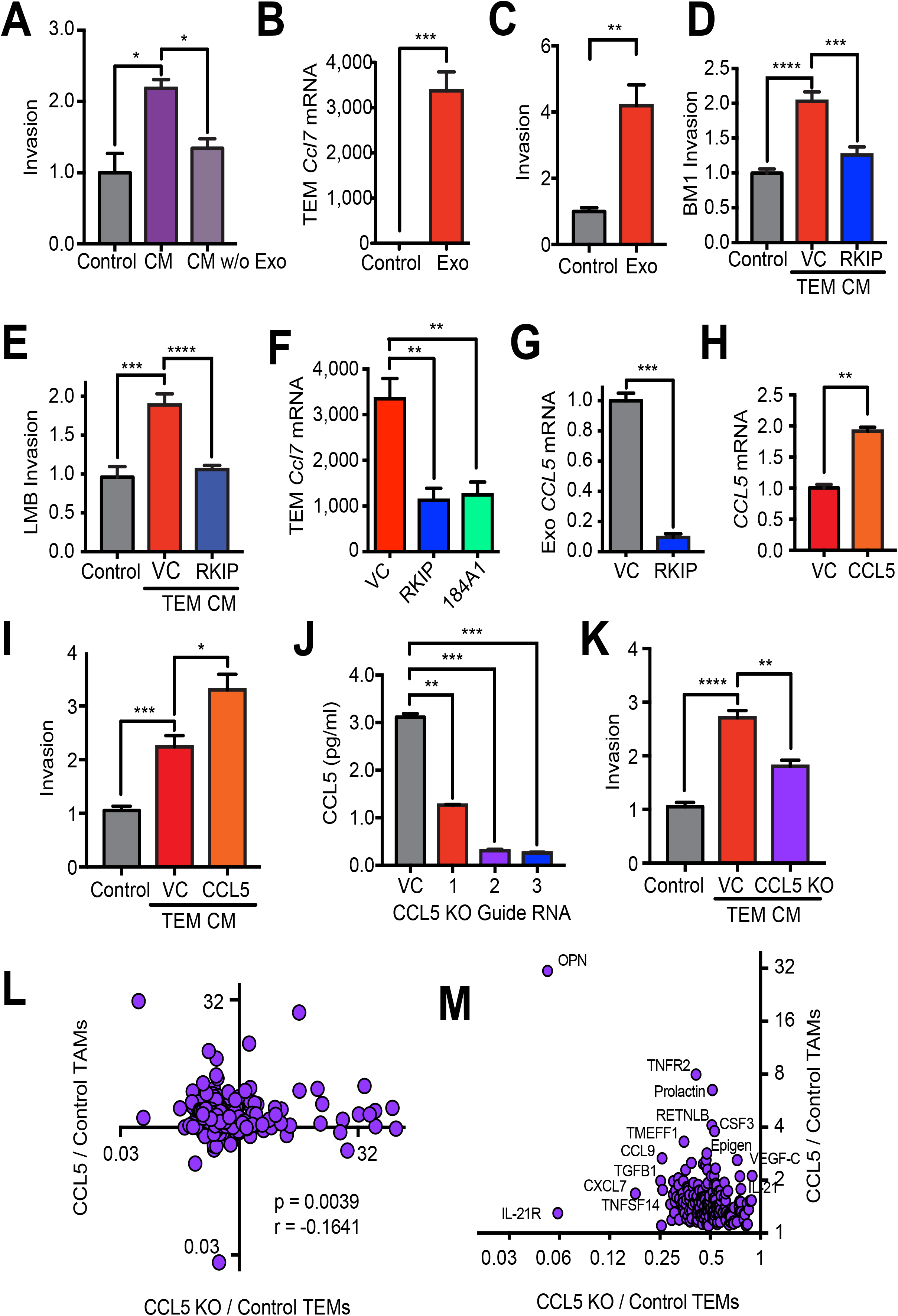
EV programming reflects the invasive state of the tumor and is regulated by CCL5. **A)** Relative invasion of BM1 cells treated with TEM CM. TEMs were programmed with M-CSF alone (Control), BM1 CM, or BM1 CM with EVs eliminated by ultracentrifugation at 100,000 x g for 1 hour (CM w/o Exo). P-values were calculated using a Student’s T-test, N=5 per group. **B)** qRT-PCR expression of Ccl7 in TEMs treated with M-CSF (Control) or M-CSF and tumor EVs (Exo) (N=4). **C)** Relative invasion of BM1 cells treated with TEM CM. TEMs were programmed with M-CSF (Control) or M-CSF and EVs (Exo). N = 5 per group. **D-E)** Relative invasion of TNBC cells treated with TEM CM. TEMs were programmed with M-CSF alone (Control), EVs from BM1+control vector (VC), BM1+RKIP, LMB+control vector (VC), or LMB+RKIP. **F)** qRT-PCR of *Ccl7* in TEMs programmed with EVs from BM1, BM1+RKIP, or 184A1 cells. **G)** qRT-PCR of *CCL5* in EVs from BM1+Control vector (VC) or BM1+RKIP (RKIP) with *GAPDH* as a loading control. **H)** qRT-PCR of *CCL5* in BM1+control vector (VC) or BM1+CCL5 (CCL5) cells using *GAPDH* as loading control (N=3). **I)** Relative invasion of BM1 cells treated with TEM CM. TEMs were programmed with CM from CSF-1 alone (Control), BM1+control vector (VC) or BM1+CCL5 (CCL5) (N=5). **J)** CCL5 protein concentration in BM1 KO cells using a Ray Biotech ELISA. *CCL5* was knocked out (KO) using a CRISPR/Cas9 lentiviral vector with indicated targeting guide RNAs for *CCL5*. CCL5 protein was measured in the CM of BM1 tumor cells collected over 24 hrs. in serum-free media and then concentrated 10X (N= 3). **K**) Invasion of BM1 cells treated with CM from TEMs. TEMs were programmed with CM from CSF-1 alone (Control), BM1+Control vector (VC), BM1+CCL5 (CCL5), or BM1+CCL5 KO (KO) (N=5). **A-K**) P-values were calculated using a Student’s t-test. **L)** Spearman correlation of cytokines secreted by TEMs or TAMs. X-axis: comparison of proteins in BM1+CCL5 KO TEMs to BM1 Control TEMs. Y-axis: comparison of proteins in BM1+CCL5 TAMs compared to BM1 Control TAMs. (N= 3 mice [TEMs] or 5 mice [TAMs], pooled). **M)** Zoom-in of the upper left quadrant in L showing proteins that are increased in TAMs by CCL5 expression and reduced in TEMs by *CCL5* KO in tumor cells.

Before directly testing the role of EVs in TEM programming, we first evaluated different methods of EV isolation to identify the one that gave the best and most consistent results. NanoSight analysis of ultracentrifuged samples, along with those isolated using ExoQuickTC reagent, showed larger aggregates relative to those isolated using the qEV size-exclusion columns (SECs) **(Fig. S1)**. EVs isolated using the qEV columns were also the most consistent and effective at programming TEMs to drive TNBC invasion **(Fig. S1)**.

Once we had established qEV SECs as the best method for isolating tumor EVs, we asked if the tumor EVs were sufficient to program TEMs. To determine if EVs in the tumor CM were responsible for TEM programming, we isolated them using qEV SECs and then added them directly to the BMDMs, as described in Fig. S1, at a concentration similar to what was found in tumor CM. The results show that EVs directly program BMDMs to induce *Ccl7* gene expression (**Fig. 3B, S2**), and EV-programmed TEM CM is sufficient to increase TNBC cell invasion (**Fig. 3C, S2**).

Our previous work established the role of metastasis suppressors such as RKIP in blocking both TAM recruitment and function (Frankenberger et al., 2015). We therefore tested whether RKIP also inhibits EV programming of pro-invasive TEMs. As anticipated, EVs from RKIP over-expressing BM1 or LMB tumor cells do not educate TEMs to potentiate tumor cell invasion (**Fig. 3D-E**). Similarly, pro-metastatic genes *Ccl7*, *Mmp12*, and *Grn* are all expressed at lower levels in TEMs programmed by BM1 RKIP over-expressing cells, comparable to levels observed in TEMs programmed by EVs from normal human mammary epithelial cells (184A1) (**Figs. 3E-F, S2**). RKIP over-expression did not significantly alter the number of EVs secreted by TNBC cells (**Fig. S2**), suggesting that EV cargo is changing. Because our previous work showed that RKIP suppresses CCL5 gene expression in the tumor cells (Frankenberger et al., 2015) and EVs can transfer mRNAs to be translated in target cells (Skog et al., 2008), we examined CCL5 mRNA levels in EVs. As in the tumor cells, CCL5 mRNA was significantly reduced in EVs from BM1+RKIP cells compared to control (**Fig. 3G**), indicating that the RNA content of the EVs is indeed altered by expression of the metastasis suppressor in tumor cells. Taken together, these results suggest that the metastatic phenotype of the tumor cell dictates EV cargo and subsequent programming of the microenvironment to reflect the invasive state of the tumor.

Because CCL5 acts on TNBC cells to potentiate TEM programming through tumor cell EVs, we asked if altering CCL5 signaling in tumor cells also impacts EV programming of TEMs. A 2-fold increase in CCL5 transcripts in tumor cells transfected with a CCL5 expression vector resulted in an ~1.5-fold increase in invasion in response to TEMs programmed by EVs from these tumor cells (**Fig. 3H-I**). These results indicate that CCL5 can regulate EV cargo in tumor cells through autocrine signaling.

Since CCL5 also regulates TAM recruitment to TNBCs (Frankenberger et al., 2015), we determined if CCL5 expression in TNBC cells is required for pro-invasive programming of TEMs by tumor EVs. Upon transduction of BM1 with a CRISPR/Cas9 lentivector and guide RNAs targeting CCL5, we efficiently knocked out (KO) the expression of CCL5 in BM1 cells (**Fig. 3J**). For subsequent experiments we utilized CCL5 KO cells generated from guide RNA 2; this construct effectively suppressed CCL5 expression but minimally affected autonomous cell growth. CCL5 KO did not change the size of EVs secreted by BM1 cells, but did yield a small but significant decrease in the number of EVs secreted **(Fig. S2)**. CM from TEMs programmed with EVs from BM1 CCL5 KO cells no longer potentiated TNBC invasion to the same degree as CM of TEMs programmed from control BM1 cells (**Fig. 3K**). These results suggest that CCL5 signaling to TNBC cells is both necessary and sufficient for maximal pro-invasive TEM programming by EVs.

To assess the physiological significance of these findings, we assayed CM from TEMs programmed by BM1 CCL5 KO or control EVs using L308 mouse cytokine arrays to identify differentially expressed cytokines. We then compared cytokines secreted by CCL5-regulated TEMs in culture to cytokines secreted by TAMs recruited to CCL5-expressing tumor cells in mice that we previously characterized (Frankenberger et al., 2015). Initially, we identified differentially expressed cytokines in the CM of TEMs programmed by control versus CCL5 KO tumor EVs using a L308 mouse cytokine array. Cytokines (62%) decreased in TEMs upon CCL5 knockout in TNBC cells overlap with cytokines previously identified as CCL5-upregulated in TAMs recruited to TNBC xenografts (Frankenberger et al., 2015a) (**Fig. 3M; see Fig. S3 for top 30**). The complete set of differentially expressed cytokines in these CCL5-downregulated TEMs versus CCL5-upregulated TAMs showed significant negative correlation (Spearman r = − 0.1641; p = 0.0039), with osteopontin (OPN) being the most strongly regulated by CCL5 in both sets (**Fig. 3L-M**). These findings further support the importance of TNBC-expressed CCL5 in regulating programming of both TEMs and TAMs to a pro-invasive phenotype through induction of pro-metastatic factors.

### EVs transmit a drug resistance phenotype from tumor cells to macrophages following treatment with the CCL5 inhibitor Maraviroc

Because of previous studies suggesting that the blockade of CCL5 could efficiently reduce TNBC metastasis (Robinson et al., 2003; Velasco-Velázquez et al., 2012) and our data showing the critical role of CCL5 in EV programming, we tested the CCR5 inhibitor Maraviroc in our system. Our previous study showed that limited Maraviroc treatment of xenograft mice could reduce tumor growth and TAM infiltration (Frankenberger et al., 2015). To determine whether Maraviroc is still inhibitory after long term treatment, we increased our treatment period to a full 21 days and began treatment earlier. However, after treating tumor-bearing mice with Maraviroc by oral gavage twice daily for 21 days, we found that tumors were significantly larger (**Fig. 4A**), and tumor cell intravasation into the blood stream actually increased (**Fig. 4B**). This tumor resistance to therapy was associated with an increased expression of CCL5 in tumor cells isolated from treated mice (**Fig. 4C**) in conjunction with higher expression of the pro-metastatic gene *Ccl7* in TAMs isolated from these mice (**Fig. 4D**).

**Figure 4:**
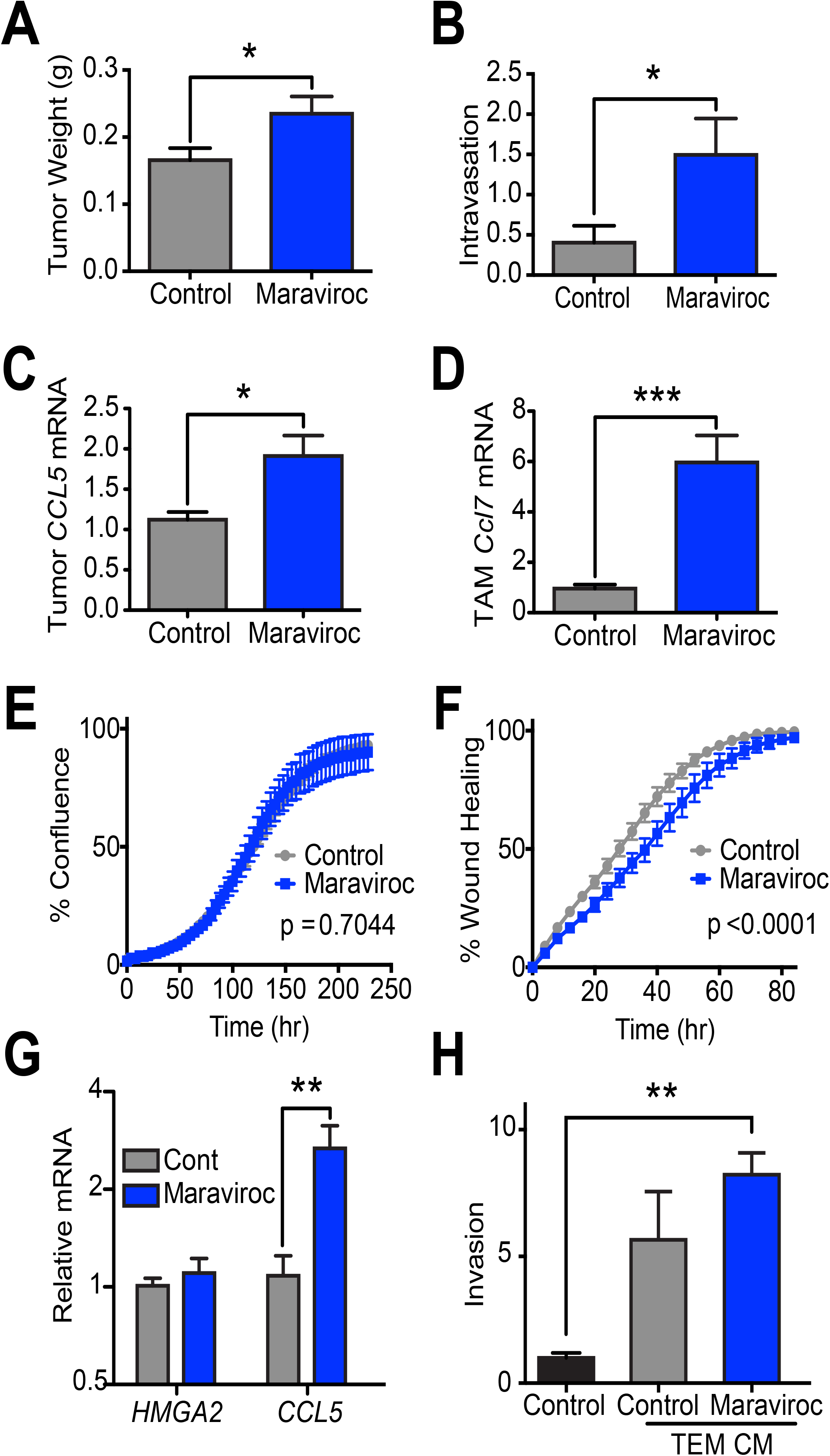
EVs transmit a drug resistance phenotype from tumor cells to macrophages following treatment with the CCL5 inhibitor Maraviroc. **A-F)** BM1 tumors from mice treated for four weeks with 8 mg/kg of Maraviroc twice daily by oral gavage **A)** Tumor weight **B)** Relative intravasation, Human *GAPDH* relative to mouse *Gapdh* from circulating cells in the blood. **C)** qRT-PCR of isolated tumor cell *CCL5* levels with *GAPDH* loading control. **D)** Mouse *Ccl7* measured by qRT-PCR relative to mouse *Gapdh* in isolated TAMs. (A-D) P values computed using a Student’s t-test **E)** % confluence. **F**) % wound healing in isolated tumor cells measured using an Incucyte every four hours. P-values calculated using a paired t-test (N=8) **G**) qRT-PCR of *CCL5* normalized to *GAPDH* from BM1 cells grown for four weeks in Maraviroc. (N=4) **H)** Relative invasion of BM1 cells treated with TEM CM. TEMs were programmed with EVs isolated from Maraviroc-treated BM1 cells. P-values were determined using a Student’s t-test (N=5).

Since EV cargo appears to be dictated by tumor cell phenotype, and, in particular, CCL5 expression, we investigated whether EVs could transmit Maraviroc resistance in tumor cells to macrophages. To generate Maracviroc-resistant cells, BM1 cells were treated *in vitro* with a low dose of drug for four weeks. Although Maraviroc did not increase growth and showed only a slight decrease in migratory ability of the cells (**Fig. 4E-F**), Maraviroc-treated tumor cells exhibited an almost 3-fold increase in the expression of CCL5 (**Fig. 4G**), similar to increases observed after *in vivo* treatment (**Fig. 4C**). When we isolated EVs from the Maraviroc -treated cells and used them to program TEMs, we found that the CM from these TEMs showed an increased ability to drive invasion in BM1 cells (**Fig. 4H**). Thus, TEMs programmed by Maraviroc-resistant, CCL5-induced cells resemble TEMs programmed by EVs from tumor cells transfected with a CCL5 expression vector. These results demonstrate that tumor EVs are capable of transmitting resistance phenotypes from tumor cells to macrophages.

### EVs are required to program TAMs to a pro-tumor phenotype

Since EVs are required for TEM programming *in vitro*, we wanted to determine if EV secretion by tumors is necessary to program TAMs toward a pro-metastatic phenotype *in vivo*. Previous work established the critical role of Rab27a in the fusion of multi-vesicular bodies to the plasma membrane prior to EV secretion (Ostrowski et al., 2010; Peinado et al., 2012), and Rab27a is highly correlated to CCL5 expression in TNBC patients (**Fig. 7B, S4**). To examine if tumor EVs are essential for programming of TAMs, we knocked out expression of Rab27a in human BM1 and mouse LMB tumor cells using lentiviral shRNAs (**Figs. 5A, S3**). We saw a corresponding decrease in EV number (down to ~50%) in the CM after transduction with multiple shRab27a clones (**Figs. 5B, S3**). When we measured EV protein, we saw a similar decrease upon Rab27a depletion (**Fig. 5C**). Since the Rab family of proteins are critical regulators of endosomal sorting and membrane fusion, we examined if Rab27a knockdown interfered with autonomous cellular growth, migration, or invasion. When Rab27a was knocked down using multiple clones of shRab27a, no change was observed in tumor cell growth, migration, or invasion **(Fig. S3)**. However, when TEMs were programmed by EVs from Rab27a-knockdown tumor cells, TEM-induced tumor cell invasion was inhibited (**Fig. 5D, S3**). These results provide additional evidence that, *in vitro,* EVs are essential for programming TEMs to potentiate tumor cell invasion.

**Figure 5:**
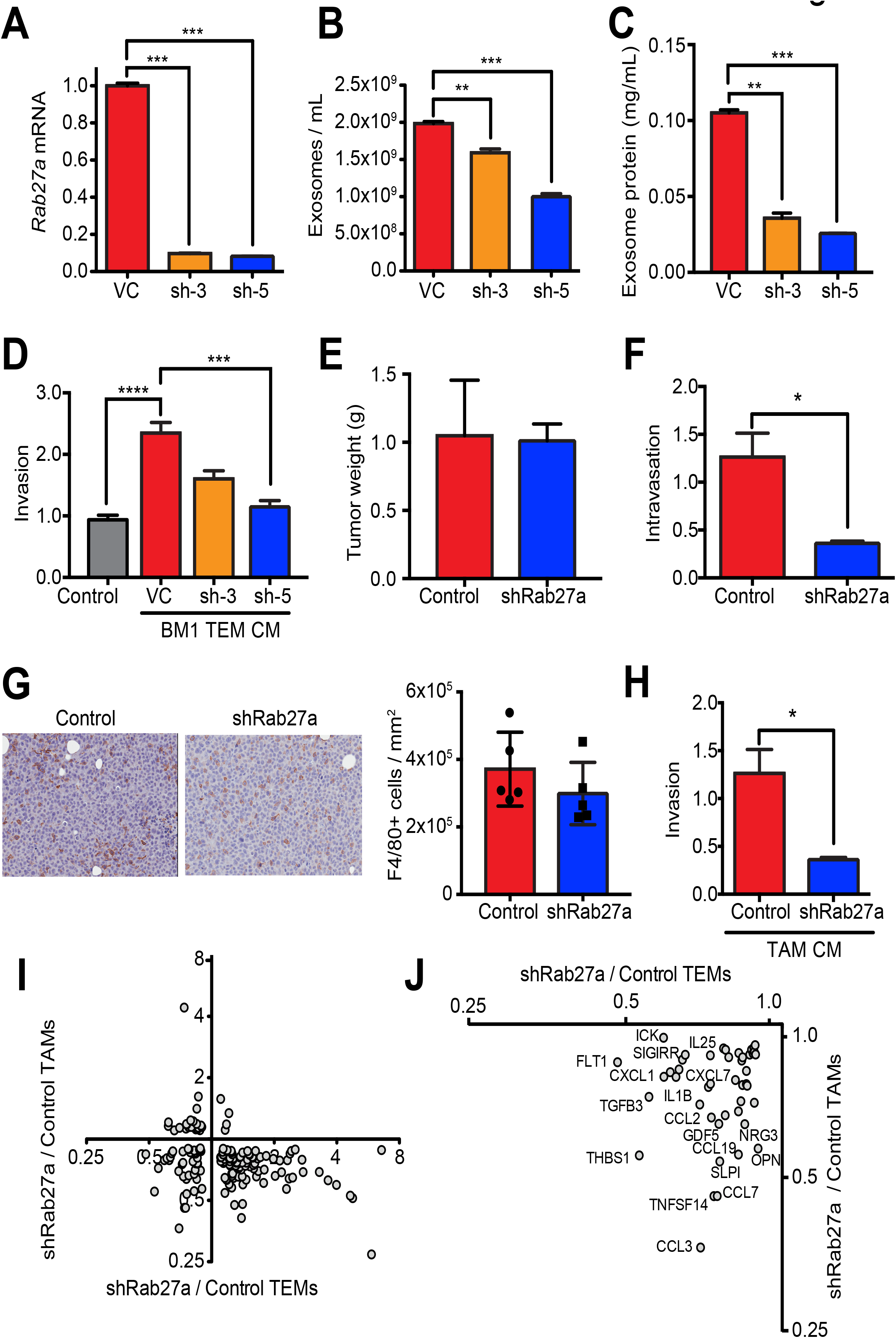
EVs are required to program TAMs to a pro-tumor phenotype. Rab27a was knocked down using a shRNA lentiviral vector, or a non-specific shRNA control. **A)** Rab27a expression measured by qRT-PCR, normalized to GAPDH expression. **B)** EV numbers measured in the CM of tumor cells after 24 hours using nanoSight. **C)** EV protein determined by pelleting EVs using ExoQuickTC from tumor CM collected in serum-free media. Protein amount was quantitated in RIPA lysates using a BCA assay. **D)** Invasion assay of BM1 cells pretreated with the indicated TEM CM. TEMs were programmed with EVs isolated from control BM1+Control vector (VC) or BM1+shRab27a (sh-5). **E-J)** BM1+Control vector (VC) or BM1+sh-5 (Rab27a) cells (2×10^6^) were injected into athymic nude mice. **E)** Tumor weights (N=5). **F)** Relative intravasation measured by quantifying the ratio of human *GAPDH* (tumor) to mouse *Gapdh* by qRT-PCR (N=5) from blood taken immediately prior to sacrifice. **G)** Representative images of F4/80 stained tumor sections in Control and shRab27a tumors. Macrophage infiltration was quantified as the number of positive cells normalized to area using Aperio. (N=5 per group). **H)** Invasion assay of BM1 cells treated with TAM CM from control or shRab27a expressing BM1 (N=5). **I)** Spearman correlation of cytokines secreted by TEMs or TAMs. X-axis: Spearman correlation of cytokines secreted by TEMs or TAMs. X-axis: comparison of proteins in shRab27a to Control TEMs. Y-axis: comparison of proteins in shRAb27a versus Control TAMs. (N= 3 mice [TEMs] or 5 mice [TAMs], pooled). **J)** Zoom-in of lower-left quadrant showing the 42 proteins reduced by shRab27a in both TEMs and TAMs. (All p-values were calculated using a Student’s t-test)

To determine if EVs are necessary for programming of TAMs *in vivo*, we chose BM1 shRab27a tumor cells with the lowest EV number (sh-5). Tumors grown with shRab27a expressed significantly less Rab27a as determined by immunoblotting **(Fig. S3)**. Knocking down Rab27a did not affect the size of BM1 tumors (**Fig. 5E**). However, it did decrease the ability of tumor cells to intravasate into blood vessels (**Fig. 5F**). As previously noted **(see Fig. S3)**, shRab27a did not affect cell autonomous invasion, suggesting that the effects on intravasation are due to changes in stromal cells within the tumor microenvironment. We therefore determined whether tumor EV secretion affected the recruitment and/or phenotype of TAMs. Rab27a knock-down did not significantly change the recruitment of TAMs to the tumor core (**Fig. 5G**). By contrast, TAMs isolated from shRab27a tumors were unable to potentiate tumor cell invasion at the same level as those from control tumors (**Fig. 5H**). These results demonstrate that blocking tumor EV secretion significantly inhibited the ability of TAMs to promote tumor cell invasion *in vitro* and tumor cell intravasation *in vivo*,

To determine the degree of functional similarity between EV-programmed TEMs and TAMs, we examined how much overlap there was between cytokines differentially expressed in both TEMs and TAMs in response to control versus shRab27a-expressing tumor cells using an L308 mouse cytokine array (**Fig. 5I**). Since EV depletion was limited to ~50%, we focused on proteins that showed relative expression levels that differed by <0.9- and >1.1-fold, resulting in 159 differentially expressed proteins. To determine relative expression of proteins between the groups, we plotted differential TEM expression along the x-axis and differential TAM expression along the y-axis. Overall, we observed that 46 of 159 differentially expressed proteins were positively correlated between the two groups (~ 30 percent overlap in differentially expressed genes within TEMs and TAMs), suggesting a significant fraction of the genes programmed in TAMs derive directly from tumor EVs. EV-regulated proteins reduced in both groups included CCL7, CCL19, CXCL1, NRG3, OPN, and TGF-beta 3 (**Figs. 5J, S3**). These proteins are also regulated by CCL5 signaling to tumor EVs **(see Fig. 3M**). These analyses suggest that tumor EVs are an essential component for programming macrophages toward a pro-metastatic TAM phenotype.

### EV-programmed TEMs promote metastasis of TNBC tumors through tumor CCL5 and TEM TLR signaling

To address the mechanism by which TNBC EVs trigger reprogramming of BMDMs to alter cytokine synthesis and secretion, we determined if TLR signaling was required. Previous work has implicated TLR signaling and NF-κB activation in tumor EV re-programming of mouse macrophage cell lines (Alexopoulou et al., 2001; Bretz et al., 2013; Chow et al., 2014). When macrophages were programmed with tumor EVs in the presence of inhibitors of NFκB, TLR1/2, or TLR3, TEM CM was no longer able to drive BM1 tumor cell invasion (**Fig. 6A**). TLR2 and TLR3 KO macrophages also lost the ability to drive BM1 invasion (**Fig. 6B**). When we compared the CM of TEMs from TLR2 and TLR3, we found a strong, positive correlation (Spearman r = 0.6681, **Fig. 6C**). TLR KO mouse TEMs compared to TEMs programmed by EVs from BM1 CCL5 KO EVs were also highly correlated (**Fig. 6D-E**). Among the cytokines identified were pro-metastatic factors such as CXCR4, CCL21, OPN, SDF1 (the ligand for CXCR4), NRG3, and TGF-beta 3 **(see Table S1)** (Hijazi et al., 1998; Moses and Barcellos-Hoff, 2011; Müller et al., 2001; Yun et al., 2011). OPN was previously shown to act as a systemic ‘instigator” to promote metastasis in breast cancer (McAllister et al., 2008). Similarly, we have shown that CXCR4 and OPN are sufficient to drive metastasis in TNBC xenografts (Yun et al., 2011). These results suggest that CCL5 signaling in the tumor and TLR signaling in the macrophages are required and act together in driving macrophages toward a pro-tumor phenotype.

**Figure 6:**
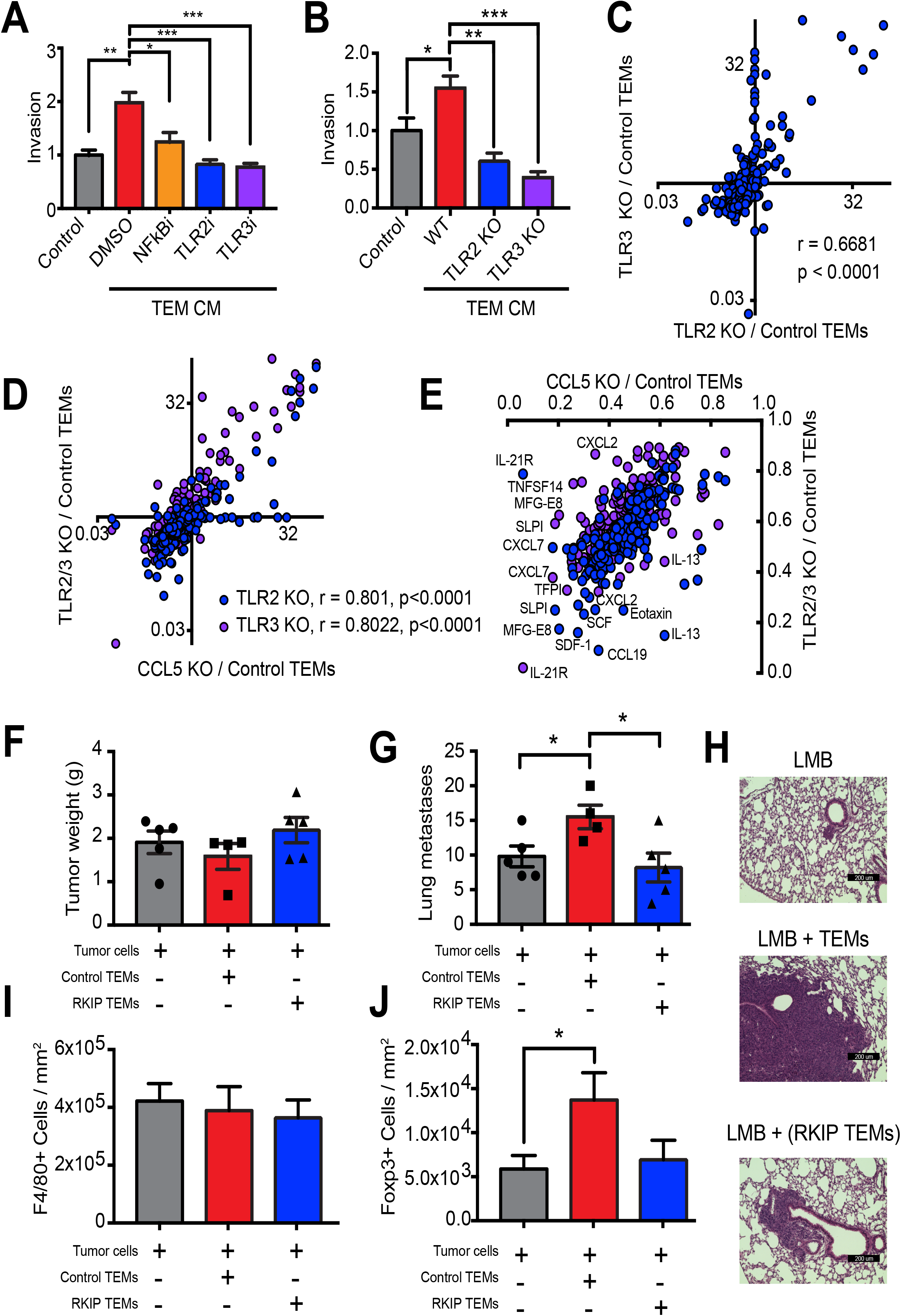
EV-programmed TEMs promote metastasis of TNBC tumors through tumor CCL5 and TEM TLR signaling. **A)** Relative invasion of BM1 cells treated with TEM CM. TEMs were programmed with BM1 EVs and inhibitors of NFkB, TLR1/2, or TLR3. (N=5) **B)** Relative invasion of BM1 cells treated with TEM CM from TLR2 and TLR3 KO TEMs. (N=5) **C)** Spearman correlation between TLR2 and TLR3. Relative protein secretion in TLR2 and TLR3 KO TEMs assayed using a mouse L308 cytokine array. (N= 3 mice, pooled) **D)** Spearman correlation between CCL5 and both TLR2 and TLR3 TEM cytokine expression. **E)** Zoom in of genes down-regulated both in TLR KO and CCL5 KO TEMs. **F-J)** Either 0.5 × 10^6^ LNB EV programmed or LNB+RKIP EV programmed TEMs were co-injected with tumor cells as indicated. **F)** Final tumor weights are shown for each mouse (N =4 or 5). **G)** Number of metastases per mouse are shown (N=4 or 5 per group). **H)** Representative images of lung metastases. **I)** F4/80+ cells were quantified using Aperio and normalized to stained area (no statistical difference) (N=4 or 5). **J)** Foxp3+ cells were quantified using Aperio and normalized to stained area (N=4). (All p-values were calculated using a Student’s t-test).

Since RKIP regulates CCL5 expression in tumor cells and EVs, we examined the effects of TEMs derived from RKIP-expressing LMB cells on metastasis *in vivo*. For this syngeneic model, we co-injected 0.5 million TEMs programmed with either LMB or LMB+RKIP EVs, with 0.5 million LMB tumor cells (LMB tumor cells alone were used as a control). Injection of TEMs with LMB tumor cells had no effect on tumor size at the end of the study (**Fig. 6F**). However, compared to LMB tumor cells alone, tumors co-injected with LMB programmed TEMs showed an increase in both number and size of lung metastases (**Fig. 6G-H**). By contrast, TEMs programmed by LMB+RKIP tumor EVs showed no significant change in lung metastases compared to control. We compared these results to the co-injection of CCL5 recruited TAMs with MDA-MB-231 tumor cells. CCL5 recruited TAMs showed an increase in both tumor growth (**Fig. 1A**) as well as tumor cell intravasation into the blood stream (**Fig. 1B**). These results suggest that TEMs, like TAMs, regulate the invasive properties of tumor cells and that EVs reflect the properties of the cells that secrete them *in vivo* as well as *in vitro*.

When we stained for F4/80 cells, we found that by the end of the study, the total number of TAMs present were the same between control and those co-injected with TEMs (**Fig. 6I**). This suggests that co-injection with TEMs does not change the ability of the tumor to recruit TAMs, but alters the phenotype of the TAMs present. Because our results strongly suggested that EV programming of TEMs drive them toward a pro-metastatic phenotype, we also immunostained tumors for Foxp3+ T-regs. Co-injection of LMB tumor cells with LMB EV-derived TEMs significantly increased the number of Foxp3+ T-regs compared to control (**Fig. 6J**). Conversely, co-injection with LMB+RKIP derived TEMs showed no change in T-reg recruitment. This result indicates that EVs, as a key regulator of macrophage phenotype, also affect the tumor microenvironment *in vivo*.

### TAM genes regulated by CCL5, TLR, and EV secretion correlate with CCL5 in human TNBC patient tumors

To determine the clinical significance of EV programming of TAMs, we examined the genes that were commonly regulated by tumor CCL5 expression, tumor EVs (shRab27a), and TAM TLR expression **(Table S1)**. DAVID (Huang et al., 2009) analysis indicates that these genes are enriched in cytokine activity, immune response, chemotaxis, cell motility and invasion, growth factor signaling, and all processes required for invasion and metastasis **(Fig. S4)**. Most of these genes are more highly expressed in TNBC or basal-like tumors compared to non-TNBC or luminal tumors (**Figs. 7A, S5**). Of these genes, we identified about half that were highly correlated with CCL5 expression across all breast tumors as a population (**Figs. 7B, S5**). When we examined the expression of these genes in TBNC tumors of individual patients, we found that these genes were also co-regulated with CCL5 in a subset of patients, particularly those categorized as ER/PR-negative (**Fig. 7C**) or Claudin-low and basal-like **(Fig. S5)**. It was noteworthy that Rab27a was among the CCL5-correlated genes, highlighting the importance of EVs in patients with CCL5 as a driver. A schematic summarizing CCL5 regulated EV stimulation of TAMs, inducing pro-metastatic cytokines through TLR signaling is shown in **Fig. 7D**.

**Figure 7:**
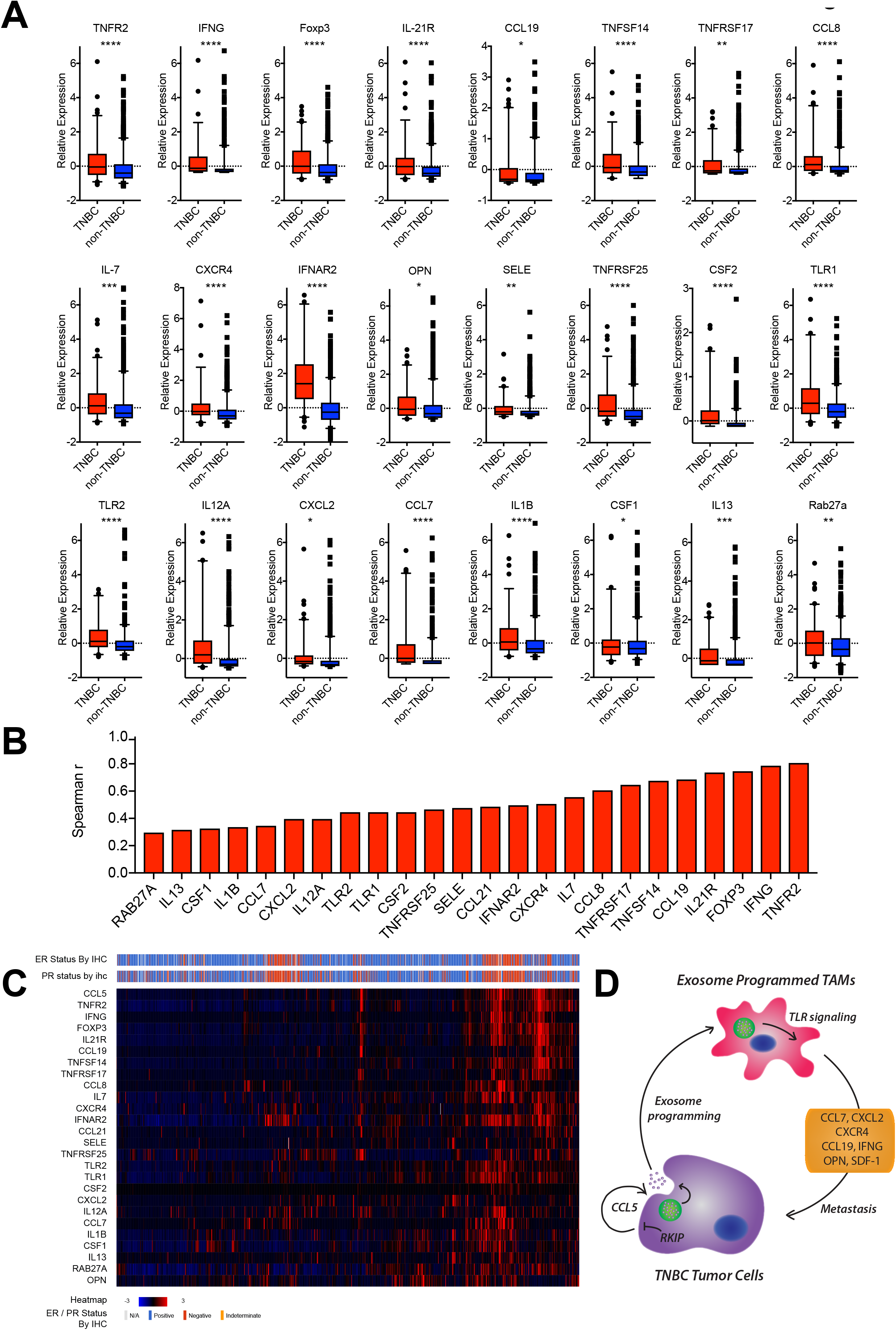
TAM genes regulated by *CCL5*, *TLR*, and EV secretion correlate with *CCL5* in human TNBC patient tumors. **A)** Normalized differential expression of genes in TNBC versus non-TNBC TCGA breast cancer samples shown as Z-scores. Genes are from the set of 52 genes regulated by EVs, tumor CCL5, and macrophage TLR signaling (**Table S1**). (TNBC, N = 103, non-TNBC = 857) **B)** Spearman correlation of genes in (**A**) to CCL5 genes in TCGA breast samples. (N = 960) **C) Heatmap showing** relative expression genes in (**A**) as well as tumor ER/PR status of TCGA breast cancer samples from individual patients. Clustering was performed using cBioPortal. (N = 960) **D)** Schematic showing programming of macrophages by tumor EVs. RKIP acts to block EV programming, while tumor CCL5 expression feeds back to increase macrophage programming. Tumor EVs regulated expression of pro-metastatic genes in TAMs and TEMs through TLR signaling to pro-metastatic cytokines. Representative cytokines are shown. (CCL7, OPN, CXCR4, SDF1, CCL19, IFNG, CXCL2.

## Discussion

Although our previous study implicated the cytokine CCL5 in the recruitment of TAMs to TNBC tumors (Frankenberger et al., 2015), the mechanism by which TAMs are educated by tumor cells was not addressed. Previous studies of the effects of tumor EVs on TAMs have relied on macrophages already programmed to a TAM phenotype. Here we demonstrate that tumor EVs alone are both necessary and sufficient for programming TAMs toward a pro-invasive phenotype in TNBC. For the fist time, we showed that blocking tumor EV secretion alone *in vivo* blocks the ability to program macrophages to a pro-metastatic, TAM phenotype. The CCL5-regulated tumor EVs induce cytokine expression in EV-educated macrophages (TEMs) through activation of TLR2 and TLR3, leading to secretion of factors that further stimulate tumor cell invasion and metastasis as well as reprogram the tumor microenvironment. We also show that altering CCL5 expression levels in tumor cells using genetic or pharmaceutical approaches regulates this process by altering EV cargo. Analyses of clinical breast cancer data showed significant correlation between expression of CCL5 and a set of cytokines in TAMs induced by the tumor EVs, and revealed specific enrichment of these cytokines in TNBC patients. Together, these results establish tumor EVs as an essential component for programming macrophages toward a pro-metastatic TAM phenotype, demonstrate that tumor EVs are capable of transmitting invasive and resistance phenotypes from tumor cells to macrophages, and provide clinical evidence supporting a role for tumor EVs in high CCL5-expressing TNBC.

The mechanism by which tumor EVs stimulate TLRs in macrophages is likely to be complex. Our results indicate that both TLR2 and TLR3 are required for programming TEMs by tumor EVs to a pro-invasive phenotype. Toll-like receptors such as TLR2 are localized on the cell surface, whereas intracellular receptors such as TLR3 reside in the endosome compartment (Kawasaki and Kawai, 2014). TLR2 was reported to bind tumor EV lipoproteins as well as miRNA (Chow et al., 2014), whereas TLR3 was reported to bind tumor EV short dsRNAs (Alexopoulou et al., 2001; Liu et al., 2016), suggesting that multiple mechanisms may be required for tumor EV programming of TEMs. The striking overlap between CCL5-regulated TEM cytokines and TLR2 or TLR3-regulated TEM cytokines provides strong evidence that both TLRs are mediators of the CCL5-regulated tumor EVs. In both these cases, the macrophages were isolated from mice. Interestingly, in human TNBC patients, CCL5 expression is most highly correlated with TLR1 and TLR2, suggesting that the specific TLRs implicated in CCL5-high expressing tumors may differ between species and possibly tumors. The nature and complexity of the cargo in TNBC EVs that trigger responses through TLRs is of significant interest but beyond the scope of the present paper.

This work also shows that CCL5 regulates macrophage recruitment and programming through two distinct mechanisms. Our previous study demonstrates that CCL5 plays a critical role in the recruitment of TAMs that drive tumor metastasis (Frankenberger et al., 2015). Since CCL5 is a well-established chemokine, CCL5 might recruit macrophages directly or indirectly via other chemokines secreted by tumor cells in response to autocrine CCL5 signaling. By contrast, the present work shows that CCL5 protein secreted from tumors does not have a direct effect on macrophage programming to a TAM phenotype. Instead, CCL5 autocrine signaling promotes generation of tumor EVs that program TEMs to a more metastatic, TAM-like phenotype. As shown directly by assay of CCL5 in EVs and indirectly through functional output, changing CCL5 expression in tumors alters EV cargo but has minimal effect on EV number. Additionally, altering EV secretion in tumors or co-injecting tumor EV-educated macrophages does not significantly change macrophage recruitment to tumors. These findings further demonstrate that CCL5 plays a novel role, through EVs, in altering macrophage phenotype rather than recruitment.

Several of the proteins that are regulated in TEMs and TAMs by tumor EVs and macrophage TLRs are potential chemotactic factors for other cell types within the microenvironment such as T-regs. In particular, CCL19 is a chemotactic factor involved in recruiting CD4+CD25+CD69-T-regs to zones of T cell infiltration in the tumor micro-environment (Guerin et al., 2011; Lim et al., 2004; Wei et al., 2006). Expression of CXCL1 also recruits T-regs to the tumor microenvironment through CXCR2 signaling (Lv et al., 2014). In addition, interferon-γ (IFNG) has been implicated in T-reg polarization (Mandai et al.; Wang et al., 2006; Wood and Sawitzki, 2006; Zaidi and Merlino, 2011). It is likely that all these factors contribute to the observed recruitment of T-regs to LMB tumors co-injected with CCL5-regulated TEMs. Since CCL19 and IFNG are highly correlated with CCL5 expression in patient breast tumors, these may be most relevant to T-reg infiltration of tumors in response to CCL5 in humans. These results indicate that tumor EVs not only impact TAM phenotypes directly, but also alter their ability to attract and interact with other cells within the tumor micro-environment. This further suggests that CCL5 and tumor EVs are key targets for tumor metastasis as well as allowing the tumor to evade immune detection.

Our work, along with other studies, suggests that EVs may also be a major mediator of drug resistance within the tumor microenvironment (André et al., 2016). For example, several studies have shown that EVs can transfer chemoresistance to recipient tumor cells through a microRNA-dependent mechanism (Chen et al., 2014; Hu et al., 2015; Patel et al., 2017). The work described here differs in that EVs transmit resistance to macrophages through a mechanism that blocks the action of a specific targeted therapy, which has not been previously demonstrated. We demonstrate that long-term treatment with the CCR5 inhibitor, Maraviroc, led to drug resistance involving up-regulation of CCL5 in tumor cells, stimulating expression of CCL7 and a pro-invasive phenotype in macrophages. Because CCL5 expression is key to EV programming of TAMs, it may be necessary to target CCL5 at the transcriptional (Ban et al., 2017) or protein assembly level rather competing for CCL5 receptor binding. Alternatively, it may be possible to target T-regs that play an essential role in blocking the anti-tumor effect of cytotoxic infiltrating CD8+ T cells through CTLA-4 and PD-1. Therefore, immunotherapy involving checkpoint inhibitors (anti-CTLA-4 and anti-PD-1/PD-L1 antibodies) might be a more effective therapeutic strategy than CCL5 inhibitors for TNBC patients with tumors that have high CCL5 expression and TAM infiltration.

Overall, our results show that tumor-associated macrophages, which play a key role in enabling tumor cells to invade, intravasate into vessels, and metastasize, rely on tumor EVs to be programmed toward a pro-metastatic phenotype. As such, EVs play an even more pivotal role in metastatic progression than previously realized. Targeting tumor EVs will not only be essential to reduce the formation of the pre-metastatic niche, but also block the programming of the primary tumor microenvironment to reduce tumor burden and tumor cell dissemination.

## Experimental Procedures

### Cell Culture

MDA-MB-436, MDA-MB-231, 293T, L929, and 4T1.2 were obtained from ATCC. MDA-MB1-231 1833 (referred to as BM1) cells were obtained from Andy Minn and E0771-LMB cells were obtained from Robin Anderson (Johnstone et al., 2015). Numerous vials were frozen upon original receipt of the cells, and all work was done within 15 passages of the initially received lines. Late passage cells were sent to Idexx for cell line authentication using STR analysis. BM1, MDA-MD-436, 4T1.2, and LMB cell lines were cultured in DMEM media supplemented with 10% fetal bovine serum, 50 U/ml penicillin, and 50 μg/ml streptomycin and L929 grown in RPMI 1640 with the same supplementation as above. Cells were transduced with lentiviral vectors for shRNA knockdown or over-expression from GE/Dharmacon. Cells were selected for 14 days using 3 μg/ml of puromycin for human cells and 10 μg/ml of puromycin for mouse cells after lentiviral transduction before use.

### Lenti-viral Transductions

All lenti-viral work was done according to institutional biosafety rules, utilizing BSL3 practices and performed in a BSL2 hood. One million 293T cells were plated in a T-25 flask the evening prior. The following day, lentiviral vectors were incubated with 3^rd^ generation viral packaging vectors and Transit LT-1 for 30 mins as described by the manufacturers protocol. DNA/LT-1 mixtures were then used to transfect 293T cells for viral production. Transfected cells were grown for 24 – 48 hours prior to viral harvesting. After incubation, viral containing media was removed, centrifuged at 2,000 x g to removed dead cells and debris, then filtered through a 0.45 um PES syringe filter. Polybrene was added to media for a final concentration of 8 ng/ml. Media was then added to target cells. Following a 24-hour transduction period, cells were washed, trypsinized and plated. All viral waste was decontaminated with 10% bleach prior to disposal. Transduced cells were then selected under high antibiotic concentrations for 10 days to ensure only transduced cells remained alive. Antibiotics were not used for subsequent cell culture.

### CCL5 Knock-Out

Lentiviral all-in-one plasmids containing Cas9 as well as guide RNAs were purchased from Applied Biological Materials. Virus was produced as described above. BM1 cells were then infected, then selected for 14 days using 3 ug/ml of puromycin. Transduced cells were assayed for CCL5 expression using a CCL5 ELISA from Ray Biotech. Samples were concentrated 10x to ensure even low levels in KO cells could be measured.

### qRT-PCR Primers

Hs GAPDH-F: TGCACCACCACCTGCTTAGC

Hs GAPDH-R: GGCATGGACTGTGGTCATGAG

Mm Hmga2-F: GAGCCCTCTCCTAAGAGACCC

Mm Hmga2-R: TTGGCCGTTTTTCTCCAATGG

Hs CCL5-F: CCAGCAGTCGTCTTTGTCAC

Hs CCL5-R: CTCTGGGTTGGCACACACTT

Mm Ccl5-F: TTTGCCTACCTCTCCCTCG

Mm Ccl5-R: CGACTGCAAGATTGGAGCACT

Mm Slpi-F: GGCCTTTTACCTTTCACGGTG Mm

Slpi-R: TACGGCATTGTGGCTTCTCAA

Mm Mmp12-F: CTGCTCCCATGAATGACAGTG

Mm Mmp12-R: AGTTGCTTCTAGCCCAAAGAAC

Mm Ccl7-F: GCTGCTTTCAGCATCCAAGTG

Mm Ccl7-R: CCAGGGACACCGACTACTG

Mm Tnfr2-F: ACACCCTACAAACCGGAACC

Mm Tnfr2-R: AGCCTTCCTGTCATAGTATTCCT

Mm Grn-F: ATGTGGGTCCTGATGAGCTG

Mm Grn-R: GCTCGTTATTCTAGGCCATGTG

Hs Grn F: ATCTTTACCGTCTCAGGGACTT

Hs Grn R: CCATCGACCATAACACAGCAC

Hs TNFR2 F: CGGGCCAACATGCAAAAGTC

Hs TNFR2 R: CAGATGCGGTTCTGTTCCC

Hs CCL5 Sequencing F: TTAGGGGATGCCCCTCAACT

Hs CCL5 Sequencing R: CTGAGACTCACACGACTGCTG

Hs Rab27a-F: GCTTTGGGAGACTCTGGT

Hs Rab27a-R: TCAATGCCCACTGTTGTGATAAA

Mm Rab27a-F: TCGGATGGAGATTACGATTACCT

Mm Rab27a-R: TTTTCCCTGAAATCAATGCCCA

### Antibodies, Cytokine Arrays, ELISAs

RKIP (derived in lab from serum of rabbits exposed to an RKIP peptide), Rabbit anti-CD63 antibody (SBI biostystem), Rab27a (AF7245, R&D Systems)

CCL5 ELISA (ELH-RANTES-1, Ray Biotech)

Mouse Cytokine Array (L308, Ray Biotech)

### Invasion Assays

As previously described, 2×10^4^ BM1 cells or 1×10^5^ E0771-LMB cells were plated in 24-well trans-well inserts with 8 μm pores (Corning) coated with growth factor depleted basement membrane extract (Trevigen)(Frankenberger et al., 2015; Surabhi et al., 2009). After incubating at 37 °C for 24 hrs, inserts were transferred to an empty well and stained with 4 ng/μl of Calcein AM (Corning) for one hour. Stained cells were gently wiped with Q-tips to remove cells on the top layer of the insert, then placed in non-enzymatic dissociation solution (Trevigen) using gentle shaking for one hour at 37 °C and 150 RMP. Fluorescence was measured using a Victor X3 fluorescent plate reader with excitation at 465 nm and emission at 535 nm.

### Tumor EV isolations

For all experiments, tumor CM was spun at 2,000 x g for 10 min to remove cell and cell debris, and then filtered through a 0.22 um PES syringe filter (Millipore). For ultracentrifugation experiments, prepared CM was spun at 100,000 x g for 70 min in polycarbonate, hard-wall tubes. EV cleared CM was then removed and saved for experiments. Remaining media was removed and EV pellets were resuspended in the same volume of serum free media from which they were isolated.

For ExoQuickTC (SBI Biosystems), prepared CM was incubated with ExoQuickTC overnight and precipitated following the manufacturers protocol.

For Izon qEV columns CM from 4 15-cm plates was prepared as above, followed by concentration using an Amicon with a 3 kD pore (Millipore) to 500 ul. Concentrated media was loaded onto a rinsed qEV column. Fractions 1-6 (column dead volume) were collected in one tume, followed by collection of EVs in fractions 7-9. Following isolation columns were cleaned with 0.5 M NaOH followed by washing with 30 mL of PBS (Corning).

### Tumor educated macrophage programming

Bone marrow was isolated from the femur and tibia of 6-10 week old C57Bl/6 mice (Charles Rivers). Red cell lysis buffer (Santa Cruz) was used to removed red blood cells. Remaining bone marrow was counted and 1 million cells were plated per well in a 6-well plate and cultured in RPMI 1640 supplemented with 10% FBS (Corning) and 50 U/ml penicillin, and 50 μg/ml streptomycin (Invitrogen) and 30% L929 Conditioned Media. Media was replenished at days two and six during culture. On day seven, bone-marrow derived macrophages (BMDMs) were washed 2-3 times with PBS (Corning) and then treated with tumor cell conditioned media or serum free media containing isolated tumor EVs supplemented with 20 ng/ml of mouse M-CSF. BMDM negative controls were grown only in serum free DMEM with 20 ng/ml of M-CSF. EV size and numbers were determined after isolation using a nanoSight microscope. 1×10^8^ EVs/mL were used for programming with BM1 tumor EVs or 1×10^9^ EVs/mL for LMB EVs.

For TEMs, after programming, cells were washed with PBS 2-3 times. Then 1 mL of serum free DMEM was added. After incubation at 37 C for 24 hours, media was removed and cells and cell debri were removed by centrifugation at 2,000 x g for 10 mins.

For co-injection, TEMs were injected into the fat pad of C57Bl/6 mice mixed with E0771-LMB cells at a ratio of 1 TEM: 1 Tumor cell. Lung metastases were assayed at the end of the study by fixing in formalin and sectioning. Six 5 um sections were quantified for number and size of metastases. Sections were 100 μm apart, and number of metastases counted were added to give number of metastases per lung.

### Tumor Associated Macrophage Isolations

As previously described (Frankenberger et al., 2015), tumors were grown to approximately 0.75 g before being harvested. Tumors were dissociated both physically with scissors to 1-2 mm pieces and using C-tubes and a gentleMACS dissociator (Miltenyi Biotech) as well as enzymatically using the human tumor dissociation kit (Miltenyi Biotech). Cells were filtered through a 70 μm mesh filter. Mononuclear cells were isolated using Ficoll-Paque PREMIUM (GE Healthcare) gradient centrifugation at 420 RPM for 45 minutes. Macrophages were then obtained using CD11b positive selection beads (Miltenyi Biotech). Flow cytometry with CD11b, F4/80, CD45, CD11c, CD205, and CCR5 was performed to determine the purity and heterogeneity of isolated TAMs.

For tumor derived macrophages, 1×10^6^ TAMs were plated in one well of a 6-well plate. After 30 minutes, cells were washed with PBS to ensure only viable macrophages attached to the plate remained. Cells were incubated for 24 hours to obtain conditioned media in serum free DMEM. Cells and cell debris were removed by centrifugation at 2,000 x g for 10 min prior to use in subsequent assays.

For co-injection studies, isolated TAMs were immediate injected into the fat pad of nude mice mixed with MDA-MB-231 tumor cells at a ratio of 1 TAM: 2 Tumor cells.

### Conditioned Media

For TEM conditioned media, cells were washed twice with PBS and once with serum free DMEM after 48 hours of programming with CM or EVs. Each well was then incubated in serum free DMEM for 24 hours to collect TEM CM. TEM CM was then spun at 2,000 x g for 10 min to remove cells and cell debris.

For tumor derived macrophages, 5×10^5^ TAMs were plated in one well of a 6-well plate. After 30 minutes, cells were washed with PBS to ensure only viable macrophages attached to the plate remained. Cells were incubated for 24 hours to obtain conditioned media in serum free DMEM. Cells and cell debris were removed by centrifugation at 2,000 x g for 10 min.

### Statistical Analysis

Statistical analysis for patient data sets was done using R and is described in those sections of Materials and Methods. Please refer there for further detail on tests used.

Otherwise, all graphics were made and all statistical analysis was done using Graph Pad Prism. Unless otherwise noted, bar graphs represent the mean (± standard error of the mean (SEM))

For comparing statistical differences between means, a T-test was used. For samples were there was no significant difference in variance between samples and samples were normally distributed or where sample size was too small to determine distribution type, a Student’s T-test was used. If sample variance differed significantly between groups, Welch’s correction was taken when the T-test was performed (does not assume equal variance between groups during test). If the samples were not normally distributed, a Mann-Whitney T test was used (does not assume normal distribution of samples).

The following was used to denote p-value unless state otherwise in Figure legends: * 0.05 > p ≥ 0.01; **0.01 > p ≥ 0.001; *** 0.001 > p ≥ 0.0001; **** 0.0001 > p

### Mice

All mice were housed and handled according to the University of Chicago Institutional Animal Care and Use Committee guidelines. Athymic nude, Balb/c mice, and C57Bl/6 were purchased from Charles Rivers. Mice were injected with 2×10^6^ human or 5×10^5^ mouse tumor cells in 100 ul of DPBS into the fatpad of 5-6 week old femal mice. Tumor volumes were measured twice per week and calculated as volume = (π/6) × width^2^ × length, where width is the shorter of the two distances. Tumors were grown to ~1,000 cm^3^ (or ~ 1 g) and removed for analysis.

Maraviroc Treatments: Tumors were allowed to engraft for 3 days prior to treatment. After 3 days, mice were treated every 12 hours for 21 days by oral gavage and tumors measured twice weekly. Control mice received 0.1 mL of water with an equivalent amount of DMSO used to dissolve Maraviroc. Treated mice received 8 mg/kg of Maraviroc every 12 hours. Statistical differences in F4/80 staining were determined by using a Mann-Whitney T-test between treated and untreated groups, while difference in tumor weight utilized an unpaired T-test with Welch’s correction. Statistical differences in tumor growth were determined using a paired T-test. A Pearson correlation was used to compare the correlation between tumor weight and %F4/80+ cells.

### Immunohistochemistry

All immunohistochemistry was performed at the University of Chicago Human Tissue Resource Center. Tissue sections were deparaffinized and rehydrated through xylenes and serial dilutions of EtOH to distilled water, and then incubated in antigen retrieval buffer at 97°C for 20 minutes. Primary and secondary antibody incubations were carried out in a humidity chamber at room temperature, and detected using an Elite kit (PK-6100, Vector Laboratories) and DAB (DAKO, K3468) system according to the manufacturers’ protocols. Following staining, tissue sections were briefly immersed in hematoxylin for counterstaining and were covered with cover glasses. Stained tissue sections were scanned at 20X magnification and analyzed using Aperio Imagescope ePathology^®^ software. To quantify infiltrating macrophages, number of F4/80 positive cells per area were measured, normalized to the area imaged.

## Author Contributions

Conceptualization, D.C.R. and M.R.R.; Methodology, D.C.R. and M.R.R.; Investigation, D.C.R., F.D.R., and J.L.; Formal Analysis, D.C.R.; Writing – Original Draft, D.C.R.; Writing – Review & Editing, D.C.R. and M.R.R.; Funding Acquisition, D.C.R. and M.R.R.; Resources, M.R.R.; Supervision, M.R.R.

## Acknowledgements

This study was funded by NIH grants R01-CA184494 (to MRR) and F31-CA192780 (to DCR) and the Rustandy fund for Innovative Cancer Research (to MRR). We are particularly grateful to Dr. Melody Swartz for generously giving us access to her NanoSight NS300 microscope. We would also like to thank members of the Swartz and Gajewsky laboratories at the University of Chicago for helpful discussions.

